# Resource use differences of two coexisting chironomid species at localized scales

**DOI:** 10.1101/2022.03.07.483209

**Authors:** Amanda R. McCormick, Joseph S. Phillips, Jamieson C. Botsch, Jón S. Ólafsson, Anthony R. Ives

## Abstract

The abundances of competing species may show positive correlations in time and space if they rely on a shared resource. Such positive correlations might obscure resource partitioning that facilitates coexistence of competitors and affects their abundances, spatial distributions, and population dynamics. Here, we examine the potential for resource partitioning between two ecologically similar midge species (Diptera: Chironomidae) in Lake Mývatn, Iceland. *Tanytarsus gracilentus* and *Chironomus islandicus* larvae coexist at high abundances in benthic habitats, and they have been previously described as feeding upon diatoms and detritus. Furthermore, both species show large, roughly synchronized population fluctuations, implying potential reliance on a shared fluctuating resource and posing the question of how these species coexist at high abundances. We first considered spatial partitioning of larvae; across multiple sites, abundances of both species were positively correlated. Thus, spatial partitioning across different sites in the lake did not appear to be strong. We then inferred differences in dietary resource use with stable carbon isotopes. *T. gracilentus* larvae had higher δ^13^C values than *C. islandicus* (mean difference = 5.39 ± 1.84‰), suggesting interspecific differences in resource use. Differences in resource selectivity, tube-building behavior, and feeding styles may facilitate resource partitioning between these two species. Relative to surface sediments, *T. gracilentus* had higher δ^13^C values (1.84 ± 0.96‰), suggesting that they selectively graze on ^13^C-enriched resources such as productive algae from the surface of their tubes. In contrast, *C. islandicus* had lower δ^13^C values than surface sediments (−2.87 ± 1.95‰), suggesting reliance on isotopically depleted resources, which may include detrital organic matter and associated microbes that larvae selectively consume from the sediment surface or within their tube walls. Overall, our study illustrates that coexisting and ecologically similar species may show positive correlations in space and time while using different resources at fine spatial scales.

## Introduction

Interactions among similar species can influence their abundances, population dynamics, and spatial distributions. For example, negative correlations among ecologically similar species are often associated with interspecific competition (Schmitt 1985, Cáceres 1998, Morin 2011), which in the most extreme cases may result in the competitive exclusion of one species from a given area (e.g., Connell 1961). Additionally, a decline in the abundance of one species (e.g., due to its specific response to an abiotic factor) may result in compensatory temporal shifts in the abundance of a competing species (Tilman 1996, Micheli et al. 1999, Klug et al. 2000). While negative correlations among species are often expected from interspecific competition, competing species may also show synchronous dynamics. Because competing species often share the same resources, resource fluctuations will affect competing species similarly, potentially leading to positive correlations in abundance (Ives et al., 1999; Ripa & Ives, 2003). Synchronous fluctuations between competitors are especially likely when their interactions with resources are strong enough to drive consumer-resource population cycles. In this case, the cyclic dynamics of one competitor will entrain the cyclic dynamics of the other, leading to positive correlations in their densities through time (Ripa and Ives 2003). Ecologically similar species may also show positive correlations across space, for example due to similar responses to the environment (Sæther et al. 2007, Kawatsu et al. 2020) or reliance on a common food source (Hellstedt et al. 2006). It follows that competitor abundances may be spatially correlated if they aggregate in areas of high shared resource abundance. Thus, even though competition tends to drive the correlation between densities of species to be more negative (Lee et al. 2020), positive associations with their shared resources might obscure the negative interactions.

Coexistence of competitors can be facilitated by mechanisms, such as resource partitioning, that weaken the strength of interspecific competition (MacArthur and Levins 1967, Pacala and Roughgarden 1982). Examples of resource partitioning include using resources at separate times (temporal partitioning), occupying distinct habitats (spatial partitioning), and specializing on different subsets of a resource, such as food (resource use partitioning) (Schoener 1974). Beyond resource partitioning, abiotic or biotic factors not directly related to resources (e.g., fluctuating environmental conditions, disturbance, predation) may promote species coexistence if they disproportionately affect a competitively superior species (Paine 1966, Leibold 1991, Gabaldon et al. 2015). Resource partitioning may be less obvious between species with similar dynamics, as reliance on a shared resource may be an integral contributor to their synchrony; yet resource partitioning may nonetheless be an important contributor to their coexistence. For example, partial partitioning of a resource (i.e., such that it is not fully shared) could be strong enough to facilitate species coexistence while still permitting enough overlap in resource use to maintain synchrony between the populations.

In this study, we compare the abundance and resource use of two ecologically similar species with synchronous population fluctuations. *Tanytarsus gracilentus* and *Chironomus islandicus* (Diptera: Chironomidae) are the predominant members of the zoobenthic community in Lake Mývatn, Iceland. As larvae, both species construct silk tubes within the sediment from which they feed. *T. gracilentus* and *C. islandicus* can occur at high abundances, with densities exceeding 330,000 and 120,000 larvae m^−2^, respectively (Ives et al. 2021). Both species show large population fluctuations, spanning five orders of magnitude for *T. gracilentus* and spanning three for *C. islandicus* (Gardarsson et al., 2004). Population crashes of *T. gracilentus* occur episodically every 4-10 years (Einarsson et al. 2002, 2004, Ives et al. 2008). Multiple lines of evidence point to consumer-resource interactions with benthic diatoms as the driver of the high-amplitude *T. gracilentus* population fluctuations (Ives et al. 2008, Einarsson et al. 2016), while other factors such as predation (Einarsson et al. 2002) or climate (Einarsson et al. 2004) appear unimportant in influencing these cycles. Furthermore, long-term monitoring of Mývatn’s chironomid populations has revealed that *C. islandicus* fluctuations are synchronous with *T. gracilentus* (Gardarsson et al., 2004), which may be due to shared reliance on similar benthic resources (Einarsson et al. 2002, Ives et al. 2008; Gardarsson et al. 2004). In support of this, previous studies suggest high dietary overlap between *T. gracilentus* and *C. islandicus*. Based on gut content analysis, late instars of both species have been described as unselectively feeding on diatoms and detritus (Ólafsson 1987 as reported within Einarsson et al. 2004); although, in studies of *T. gracilentus* gut contents, moderate variation in selectivity for diatoms and detritus was found across larval instars and seasons (Ingvason et al. 2002, 2004). Fatty acid analysis has also supported *T. gracilentus* reliance on diatoms, as well as bacteria that are likely associated with detritus (Ingvason 2002); there are no corresponding fatty acid profiles of *C. islandicus* from Mývatn. Overall, potentially high overlap in *T. gracilentus* and *C. islandicus* diets suggests that similar resource requirements may underlie their high-amplitude and synchronized population fluctuations in Mývatn.

Our objective was to consider factors that may facilitate the high abundance of both *T. gracilentus* and *C. islandicus* in the benthos of Mývatn despite the potential for competition. Specifically, we examined patterns of larval abundance and stable carbon isotopes to infer potential differences in their use of spatial and dietary resources, respectively. Although *T. gracilentus* and *C. islandicus* populations fluctuate synchronously across generations (Gardarsson et al., 2004), these long-term data are based on the counts of emergent adults and integrate midge abundances over much of the lake. This leaves the possibility that *T. gracilentus* and *C. islandicus* larvae occur in different locations or microhabitats within the lake. Therefore, we considered the potential for spatial partitioning by examining the correlation in larval abundances of *T. gracilentus* and *C. islandicus* across several locations. Furthermore, we assessed differences in dietary resource use with stable carbon isotopes, which provide information about assimilated resources because there is relatively little difference in the ratio of ^13^C/^12^C isotopes between a consumer and its diet (Rounick and Winterbourn 1986). Interspecific comparisons of δ^13^C signatures could reveal differences in resource use between *T. gracilentus* and *C. islandicus* that are not apparent from gut contents, as stable isotope signatures integrate consumer resource use over a longer period of time (Rounick and Winterbourn 1986). To identify interspecific differences in resource selectivity, we compared δ^13^C values of *T. gracilentus* and *C. islandicus* larvae to those of surface sediment, which is a putative resource for the two species. Finally, we examined sediment δ^13^C values at different depths within the sediment to consider how vertical patterns of organic matter isotope signatures may contribute to interspecific differences in larval signatures.

## Methods

### Study system

Lake Mývatn is located in northeastern Iceland (65°37’N, 17°00’W) and has high primary production, with limited allochthonous inputs (Jónasson 1979). The lake is shallow, with a mean depth of 2.3 m and maximum depth of 4.2 m in the main south basin (Jónasson 1979). The smaller north basin was historically shallower (mean natural depth 1 m), but diatomite mining from 1967-2004 created depressions >5 m deep (Fig. 1; Einarsson et al. 2004). Substrate type is the primary feature for categorizing benthic habitat as littoral or profundal in Mývatn, in contrast to other limnological applications that may base this designation on light penetration and/or depth. Benthic substrate in the nearshore littoral habitat includes rocks and sand (Lindegaard and Jónasson 1979). Mývatn’s profundal habitats are characterized by extensive soft-bottomed mud flats, consisting of sediment composed of diatom frustules, organic matter, sand, and tephra (Jónasson 1979). Benthic algae (mainly epipelic diatoms on the sediment surface) comprise a majority of whole-lake production, although phytoplankton production varies spatially and temporally within and across years, especially due to cyanobacteria blooms (Einarsson et al. 2004, Phillips 2020, McCormick et al. 2021). Available food sources for sediment-dwelling primary consumers in profundal habitats are likely limited to benthic algae, settled phytoplankton, autochthonous detritus (potentially from benthic and pelagic sources), and sediment-associated microorganisms.

**Figure 1.**
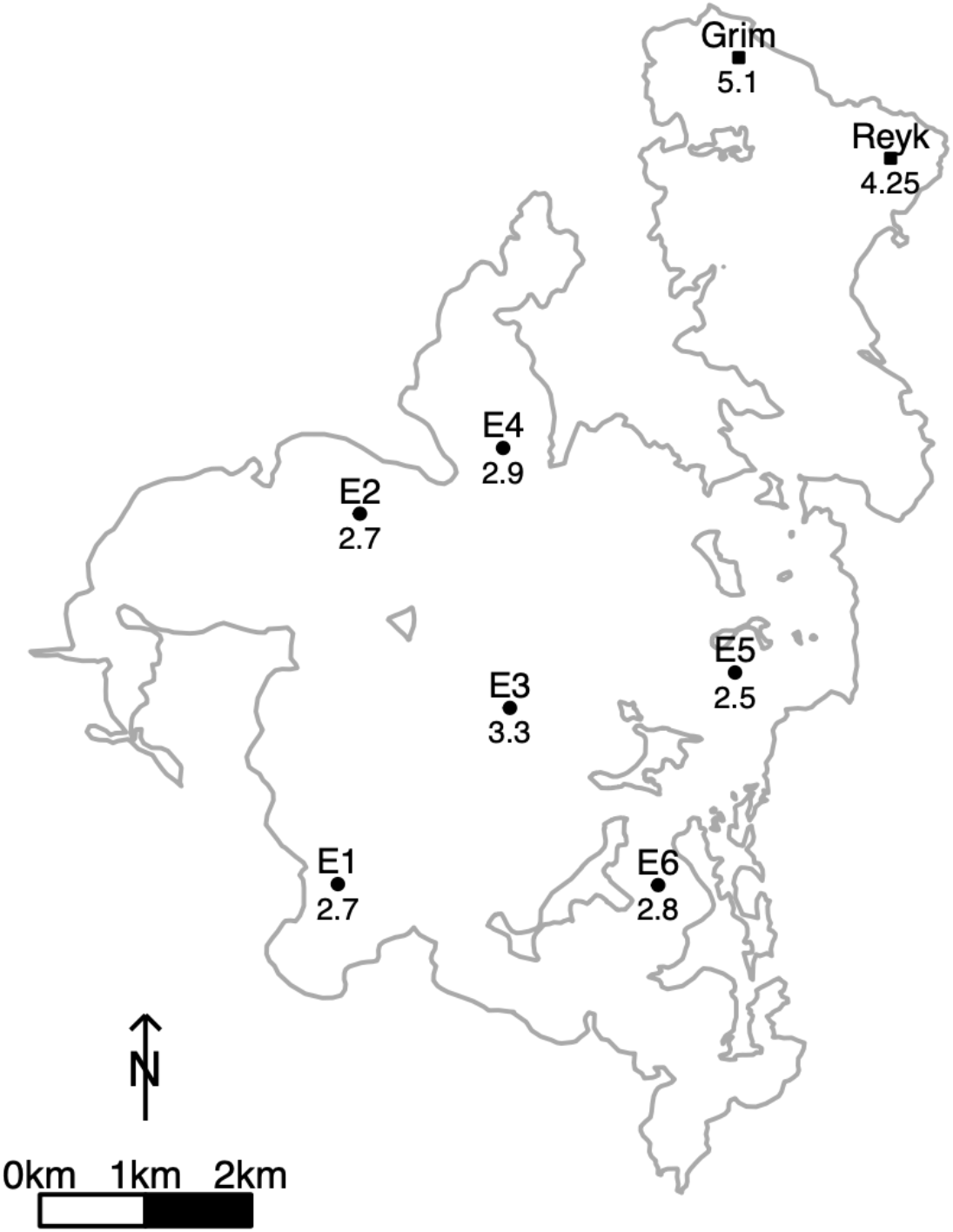
A map of Lake Mývatn depicts site locations from which surface sediment and larval midges were collected for stable carbon isotope analysis. Site names appear above each symbol, with corresponding water column depths (in meters) below. For details regarding collection dates for stable isotope samples, see Supporting Information (Table S1). Routine monitoring data from 2015-2019 from the six sites in the south basin (circles; E1-E6) were used to examine correlations in abundances of *T. gracilentus* and *C. islandicus* larvae.

Benthic pathways dominate Mývatn’s energy flow, and midge larvae are the lake’s dominant consumers in both number and biomass (Jónasson 1979). The dominant midge species in areas of sediment substrate (i.e., profundal habitat) are *T. gracilentus* and *C. islandicus* (Lindegaard and Jónasson 1979). These species are ecologically similar and reside in tubes which they construct from silk. However, the depths of their tubes differ, with *C. islandicus* burrowing deeper within the sediment than *T. gracilentus* (Herren et al. 2017). *C. islandicus* larvae are roughly an order of magnitude times the mass of *T. gracilentus*, although body size ratios vary among years and larval stages (Lindegaard and Jónasson 1979, Herren et al. 2017). Larval densities of *T. gracilentus* and *C. islandicus* may exceed 330,000 and 120,000 individuals m^-2^, respectively (Ives et al. 2021). Despite the typically lower densities of *C. islandicus* compared to *T. gracilentus*, both species contribute strongly to secondary production in years of high midge abundance, due to the higher individual biomass of *C. islandicus* (Lindegaard and Jónasson 1979). While *T. gracilentus* and *C. islandicus* both undergo larval development through four instars, an aquatic pupal stage, and adult emergence from the lake, there are notable differences in their life histories. *T. gracilentus* generally has two adult emergences per year in spring (early June) and summer (late July to early August). *C. islandicus* individuals spend either one or two years as larvae (Lindegaard and Jónasson 1979), and the annual *C. islandicus* emergence generally overlaps with the spring *T. gracilentus* emergence. For the *T. gracilentus* generation that emerges in summer, rapid larval growth in terms of ash free dry weight gain occurs in a roughly two-month summer period (Lindegaard and Jónasson 1979). For the generation that emerges in spring, rapid larval growth occurs from August to October, and then resumes the following April with the transition from third to fourth instar. Individual growth of *C. islandicus* larvae is greatest from June to October (Lindegaard and Jónasson 1979). Under a two-year life cycle, *C. islandicus* larvae spend the first winter in the third instar and the second winter in the fourth instar, while larvae that develop in one year spend a majority of time in the fourth instar, including the overwinter period (Lindegaard and Jónasson 1979). Thus, despite the life history differences between *T. gracilentus* and *C. islandicus*, their growth patterns largely overlap temporally, suggesting the importance of resources available in summer.

### Larval abundance

We routinely sampled midge larvae at multiple sites in Mývatn as part of a long-term monitoring project. From 2015-2019, larvae were sampled from six sites (Fig. 1) 3-4 times per year (from late May to mid-August). Sediment cores were collected with a Kajak corer, with replicate (n = 3-5, but typically n = 3) cores for each sampling event. The spatial extent of a single core is 20 cm^2^ (diameter = 5 cm), and replicate cores within a site encompass a few square meters. The top 1.5-cm sediment layer was extruded from each core and sieved through 63-µm mesh, and the remaining sediment was sieved through 125-µm mesh. Finer mesh was used for the top sediment to capture first instar *T. gracilentus* larvae which reside in this layer. After recombining the material retained on the 63-µm and 125-µm mesh sieves for a given core, samples were subsampled to a target of 50-100 larvae, and midges were hand-picked and stored in 70% ethanol. Larvae were identified to the tribe level (Tanytarsini and Chironomini, to which *T. gracilentus* and *C. islandicus* respectively belong) using a dissecting microscope.

While other species of Tanytarsini and Chironomini are found in Mývatn, the locations of our sites make it likely that almost all Tanytarsini were *T. gracilentus* and almost all Chironomini were *C. islandicus*. Specifically, all sites in this study were profundal, where *T. gracilentus* comprises the majority of Tanytarsini. The other species of this tribe (e.g., *Micropsectra* spp.) occur mainly in Mývatn’s littoral habitats (Gardarsson et al., 2004; Lindegaard & Jónasson, 1979). Likewise, *C. islandicus* comprise the majority of Chironomini, but other rare species in the *Chironomus* genus occur in the lake, mainly in the north basin (Lindegaard and Jónasson 1979, Einarsson et al. 2004); even so, *C. islandicus* accounts for 97.4-98.8% of the genus’ abundance in the north basin (Lindegaard and Jónasson 1979). Moreover, in samples of adult midges collected with emergence traps at the six monitoring sites from 2015-2019, all Tanytarsini were identified as *T. gracilentus* and a majority (96%) of Chironomini were identified as *C. islandicus* (Botsch, unpublished data). Because *T. gracilentus* and *C. islandicus* are dominant in the areas we sampled, they likely encompass most specimens in our study (i.e., in the larval abundance and stable isotope samples). Thus, we use these species names throughout the text; however, we acknowledge the possibility that our samples may have included a small number of individuals of other species in these tribes.

### Stable isotope samples

During the summers of 2016-2018, we collected midge larvae and surface sediment for stable isotope analysis from sites with water column depths ranging from 2.5 to 5.1 m (Fig. 1). In 2016, collection of stable isotope samples focused on the six sites from which larval abundance data were collected (see above), but in subsequent years isotope sample collection focused on the main long-term monitoring station at Mývatn (site “E3”, depth = 3.3 m) and sites in the lake’s north basin (Fig. 1). A complete list of stable isotope samples, including collection dates and locations, is in the Supporting Information (Table S1).

We obtained midge larvae for stable isotope analysis using Kajak cores separate from those used to monitor larval abundances. We sieved these cores through 125-µm or 250-µm mesh and picked individuals from the remaining material into untreated groundwater sourced from a laboratory tap, where they stayed for 48 h at 4 °C to clear their guts. We distinguished larvae as Tanytarsini or Chironomini while alive, as fixing them with a preservative (e.g., ethanol) to identify them formally would potentially affect their stable isotope values. We aimed to collect third and fourth instar individuals, which are more easily identified from live specimens than are the smaller instars. Larvae were pooled for stable isotope analysis, with the number pooled dependent on individual mass and availability. Samples included a minimum of 8 *T. gracilentus* and 2 *C. islandicus* individuals, but we aimed to use more individuals (up to ~100 *T. gracilentus* and ~30 *C. islandicus*). In general, larvae from replicate cores were combined from a site-date combination, in part to reach the necessary biomass for isotopic analysis, and also to streamline sorting in the lab; however, larvae from replicate cores were analyzed separately in some instances. Pooled larvae for stable isotope samples were dried at 60 °C.

To obtain surface sediment for stable isotope analysis, we sampled the top 0.75 cm of sediment from additional Kajak cores to capture the most photosynthetically active layer of benthic algae. These samples do not represent a pure sample of algal material and also contain surface sediment microbial assemblages and detritus; nonetheless, similar sampling techniques have been used to characterize benthic resources in other studies (Karlsson and Byström 2005). To remove midges and their tubes from sediment samples, we gently sieved material through 500-µm mesh and collected the flow-through on a 20-µm screen. We removed silken midge tubes to limit the influence of larval isotope values (i.e., from silk they produced) on the surface sediment signatures, although we acknowledge that small silk fragments may have passed through the 500-µm mesh. The retained material on the 20-µm mesh was transferred to a tin weigh boat for drying at 60 °C.

In 2019, we collected additional cores to characterize δ^13^C values at different layers within the sediment. Three cores were collected from two of the sites previously sampled for sediment isotope samples (with water column depths of 2.5 m and 4.25 m). A 0.75-cm layer of sediment was collected from the surface of these cores (as above), as well as 0.75-cm thick layers that were collected 5 cm and 10 cm below the sediment surface. These sediment samples were then processed using the procedure described above.

Dried larvae and sediment samples for stable isotope analysis were homogenized using plastic pestles, placed in a desiccator at room temperature for 24 h, and weighed into tin capsules. Analyses of stable carbon isotope ratios were conducted by the University of California Davis Stable Isotope Facility (Davis, CA, USA). Isotope ratios are expressed using delta (δ) notation (units ‰), where δ = [(R _sample_ − R _standard_)/R _standard_] × 1000, and R = ^13^C/^12^C. The reference material for carbon was Vienna Pee Dee Belemnite. Samples were run by this facility in multiple years, with the analytical errors (reported as standard deviations) for δ^13^C ranging from 0.02-0.09‰. To quantify error associated with sample preparation, duplicates were analyzed for a subset of samples, which produced standard deviations for δ^13^C of 0.029‰ for sediment (n = 3 pairs) and 0.189‰ for midge larvae (n = 5 pairs).

To assess the influence of inorganic carbonates on sediment δ^13^C signatures, we performed an HCl acidification following the methods of Harris et al. (2001) on a subset of sediment samples collected in 2018 and compared the isotopic signatures of acid-fumigated samples to non-treated samples from the same core. There was no difference in δ^13^C between acid-fumigated and non-treated sediment (paired t-test: t_7_ = −1.25, p = 0.252). Thus, we present δ^13^C signatures from non-acidified sediment samples.

### Statistical analysis

We performed analyses to examine patterns in the abundances of *T. gracilentus* and *C. islandicus* larvae and analyses comparing their δ^13^C values. We calculated the correlation in larval abundances of *T. gracilentus* and *C. islandicus* to examine whether a high abundance of one species was positively or negatively associated with the abundance of the other. For this analysis, we used the five available years of data on larval abundances within sediment cores from the six routine monitoring sites (Fig 1). We conducted the correlation in larval abundances assuming that abundances followed a lognormal-Poisson distribution (i.e., a Poisson distribution whose parameter λ is lognormally distributed) with the pglmm() function in the ‘phyr’ package (Ives et al. 2020), allowing for different lognormal variances for each species. We conducted supplemental analyses that examine the correlation in larval abundances at two aggregated levels: site-date and site-year combinations (Supporting Information).

To investigate differences among sediment and larval δ^13^C signatures, we used the mean value for date-site combinations of each sample type (i.e., using an aggregated value when replicates were available). We performed three paired analyses to compare δ^13^C values between (i) *T. gracilentus* and *C. islandicus* larvae, (ii) *T. gracilentus* larvae and surface sediment, and (iii) *C. islandicus* larvae and surface sediment. For these analyses, we were interested in paired differences between samples collected at the same time from the same location. In most cases, samples of both types from a given site were collected on the same date, but for cases in which the dates did not match exactly we paired sample types as closely in time as we could from the same site. Generally, this was a difference of 1-5 d, but in 2016 five *T. gracilentus* samples and six *C. islandicus* samples collected in late May were paired with sediment collected two weeks later (Supporting Information: Fig. S1). The full set of three sample types was not always available for a given site within a two-week period, so for each pair of sample types we analyzed the subset of our data that included paired samples for each comparison of interest. For each analysis, we investigated differences in δ^13^C values using paired t-tests and report the differences in isotope values for each sample pair.

We explored spatial variation of δ^13^C values (i) horizontally across sites in the lake as a function of the depth of the water column and (ii) vertically within the sediment profile. First, we examined how water column depth of the sample sites affected δ^13^C values of midge larvae and surface sediment with a linear model. In this analysis, we included all data for each sample type for all sampled site-date combinations from 2016-2018 (i.e., regardless of whether it was part of an above paired analysis). Second, using the cores collected in 2019, we examined how layer depth within the sediment affected δ^13^C values with a linear mixed effect model (LMM), with water column depth of the site and layer depth within the sediment as fixed effects and sediment core identity as a random effect. The LMM was fit using the ‘lme4’ package (Bates et al. 2015), and statistical results are reported from an F-test with Kenward-Roger denominator degrees of freedom using the ‘car’ package (Fox and Weisberg 2019). All statistical analyses were performed in R version 4.0.3 (R Core Team, 2020).

## Results

### Larval abundance

Combining the years 2015-2019, we analyzed the correlation among *T. gracilentus* and *C. islandicus* larval abundances using a total of 301 Kajak core samples that were taken from six sites, with each site sampled on 16-17 different dates. At the level of individual cores, *T. gracilentus* and *C. islandicus* larval abundances were positively correlated (*r* = 0.52, *P* ≪ 0.001; Fig. 2), with the lognormal variance of *T. gracilentus* (4.64) greater than that of *C. islandicus* (2.28). Abundances of the two species were also positively correlated at the levels of aggregated site-date combinations and aggregated site-year combinations (Supporting Information; Fig. S2).

**Figure 2.**
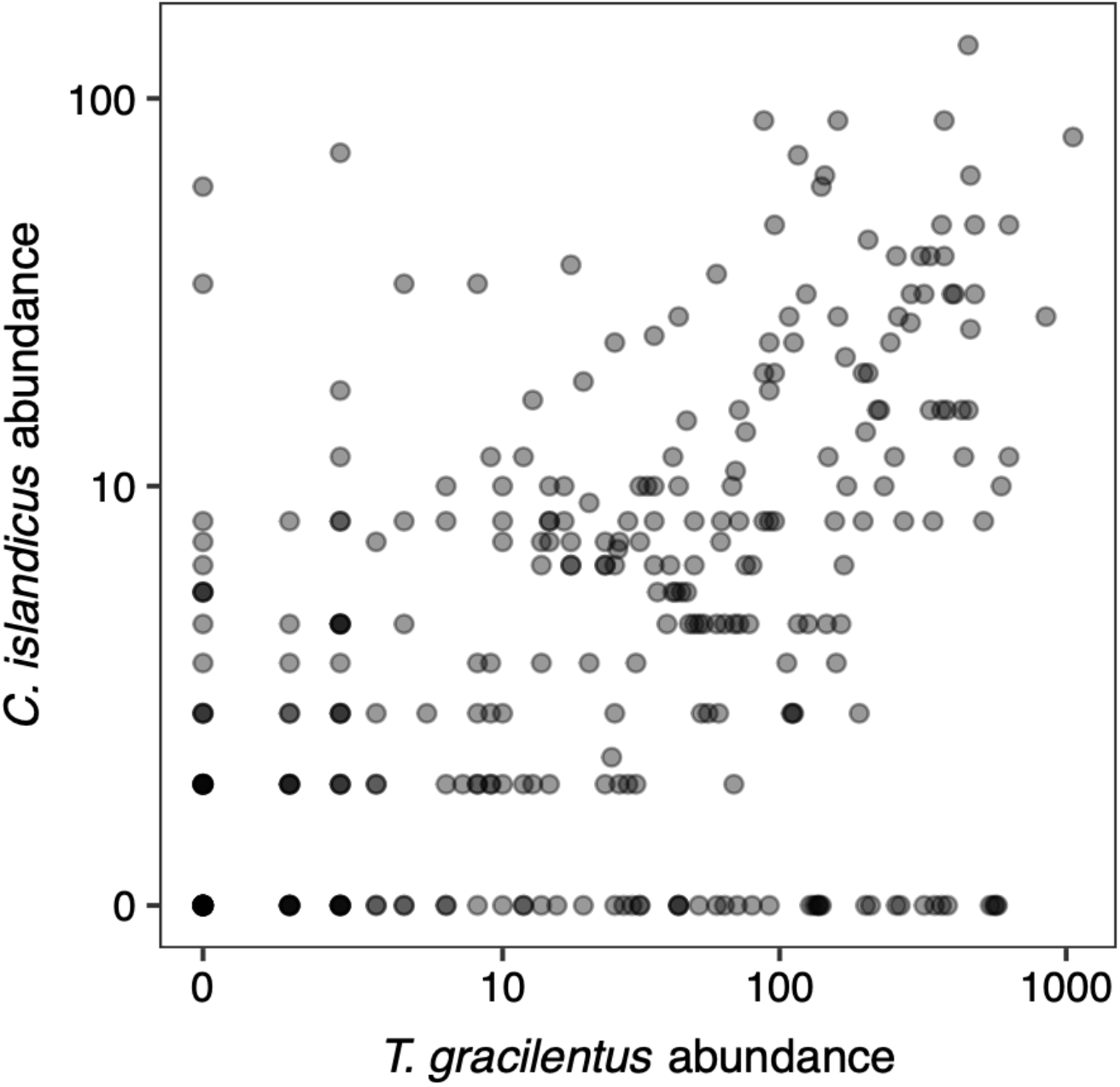
*T. gracilentus* and *C. islandicus* larval abundances in sediment cores from six routine monitoring sites in Mývatn’s south basin collected during 2015-2019. Larval abundances are shown on a log(x + 1) scale to include counts that were zero.

### Stable isotope values

Paired *T. gracilentus* and *C. islandicus* larval samples significantly differed in their δ^13^C signatures (t_11_ = 10.18, p < 0.001). *T. gracilentus* larvae consistently had higher δ^13^C values than *C. islandicus* (Fig. 3a), with a mean difference (± SD) in their δ^13^C signatures of 5.39 ± 1.84‰. Relative to paired surface sediment samples, *T. gracilentus* larvae had significantly higher δ^13^C values (t_9_ = 6.07, p < 0.001; Fig. 3b), with a mean difference of 1.84 ± 0.96‰. In contrast, *C. islandicus* larvae had significantly lower δ^13^C values compared to paired surface sediments (t_22_ = −7.07, p < 0.001; Fig. 3c), with a mean difference of −2.87 ± 1.95‰.

**Figure 3.**
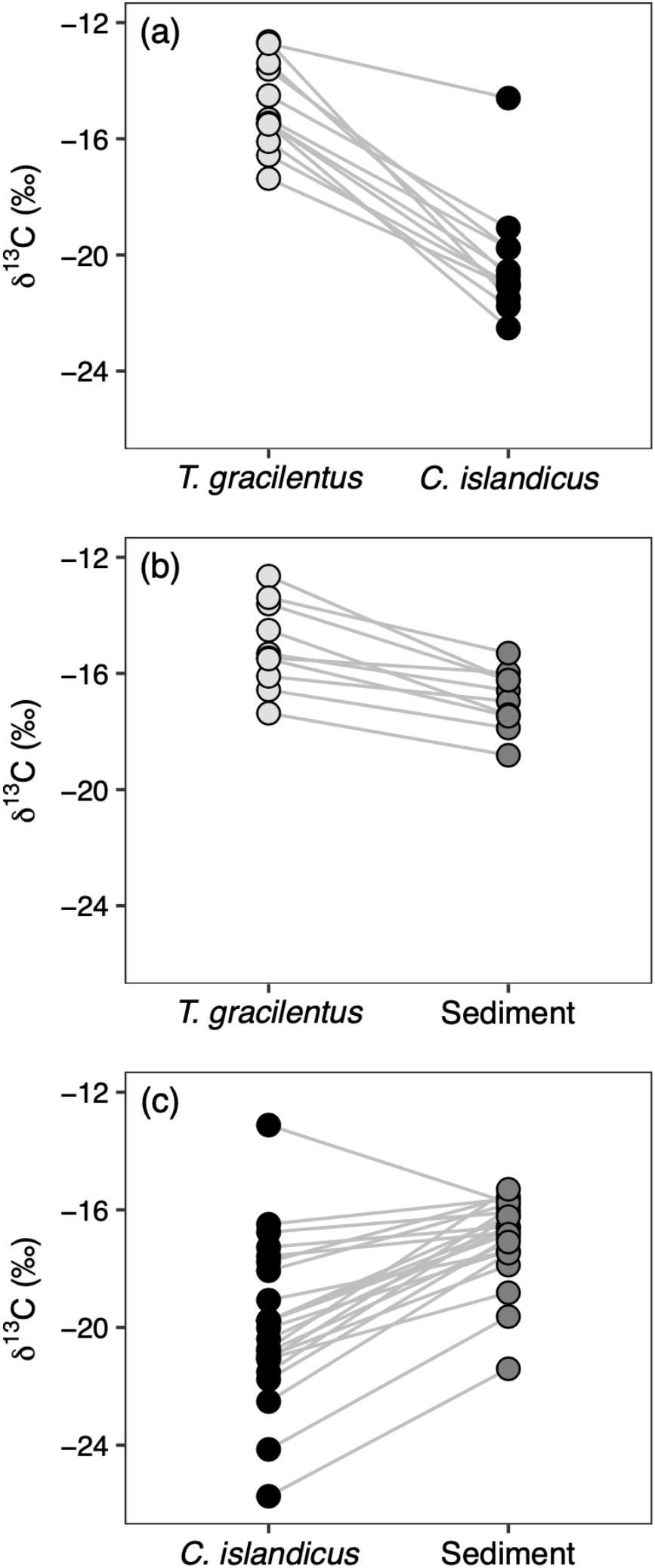
Comparisons of δ^13^C signatures among paired (a) *T. gracilentus* and *C. islandicus* larvae, (b) *T. gracilentus* larvae and surface sediment, and (c) *C. islandicus* larvae and surface sediment. Lines connect samples from a specific date-site combination that were compared to one another in each paired analysis.

*T. gracilentus* larvae, *C. islandicus* larvae, and surface sediment δ^13^C values decreased with increasing water column depth (Fig. 4; F_1, 62_ = 60.73, p < 0.001). The sample type × site depth interaction (F_2, 62_ = 3.69, p = 0.031) implied *C. islandicus* δ^13^C signatures decreased somewhat more strongly with water column depth than *T. gracilentus* or sediment. However, the slope coefficients [95% confidence intervals] for the three sample types (*T. gracilentus*: −1.52 [−2.84, −0.21]; *C. islandicus*: −3.24 [−4.08, −2.41]; surface sediment: −1.96 [−2.68, −1.24]) were close to each other (Fig. 4), reflecting the marginal statistical significance of the interaction.

**Figure 4.**
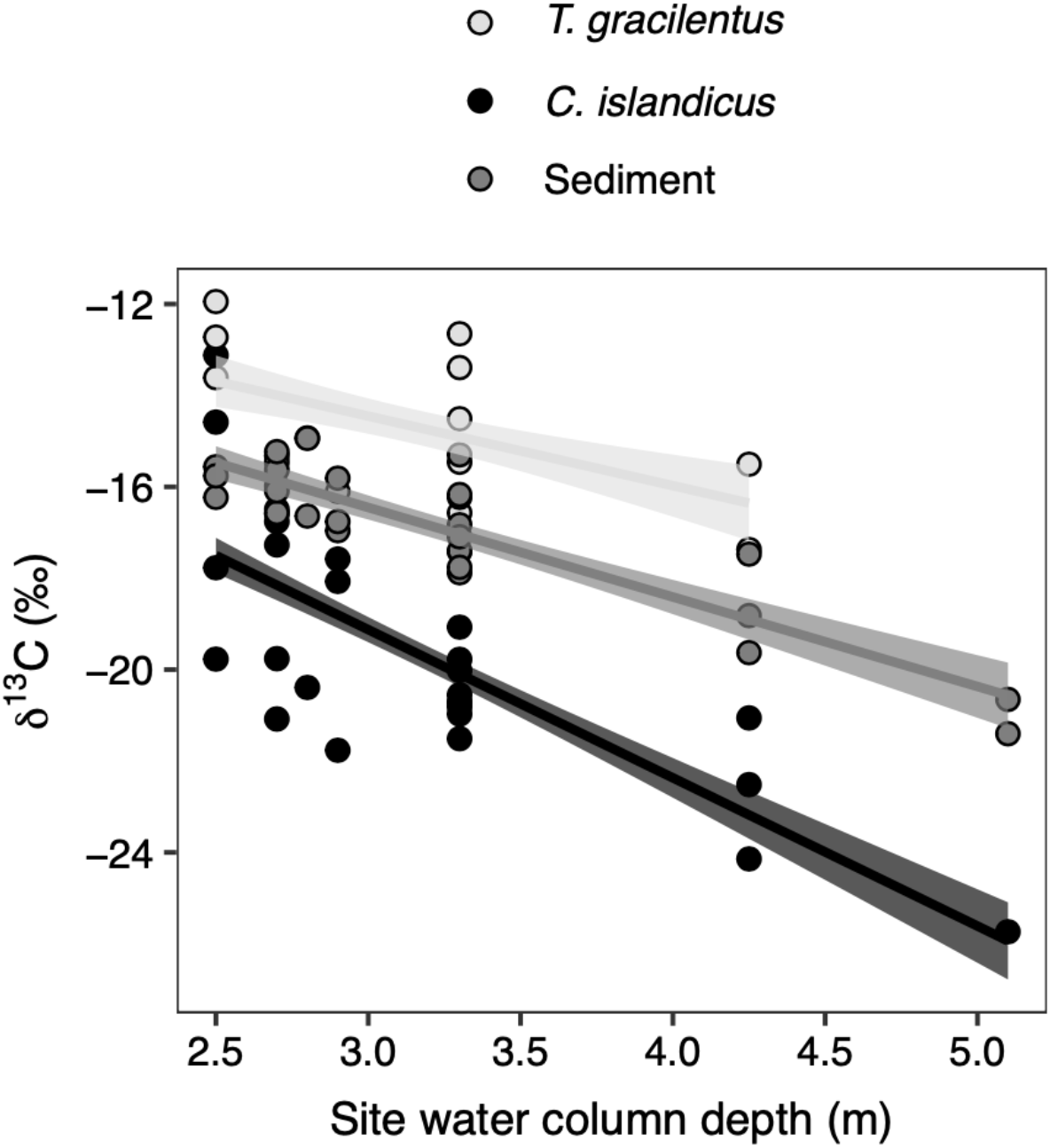
The δ^13^C signatures of *T. gracilentus* larvae, *C. islandicus* larvae, and surface sediments across sites varying in water column depth. Points show raw data, and lines correspond to the linear model fit with standard errors.

Within the sediment, layer depth significantly affected δ^13^C values (F_1, 11_ = 11.64, p = 0.006). Within a sediment core, δ^13^C values were higher in the top 0.75-cm layer than those from 5 cm or 10 cm below the sediment surface (Fig. 5). These results demonstrate vertical patterns of sediment δ^13^C values within the benthic habitat. The water column depth of the sites from which cores were collected also affected sediment δ^13^C values (F_1, 4_ = 85.34, p = 0.001). Sediment from the site with 4.25-m water depth had significantly lower δ^13^C values than the site at 2.5-m depth (Fig. 5), mirroring our results from the analysis of water column depths and sample δ^13^C values (Fig. 4).

**Figure 5.**
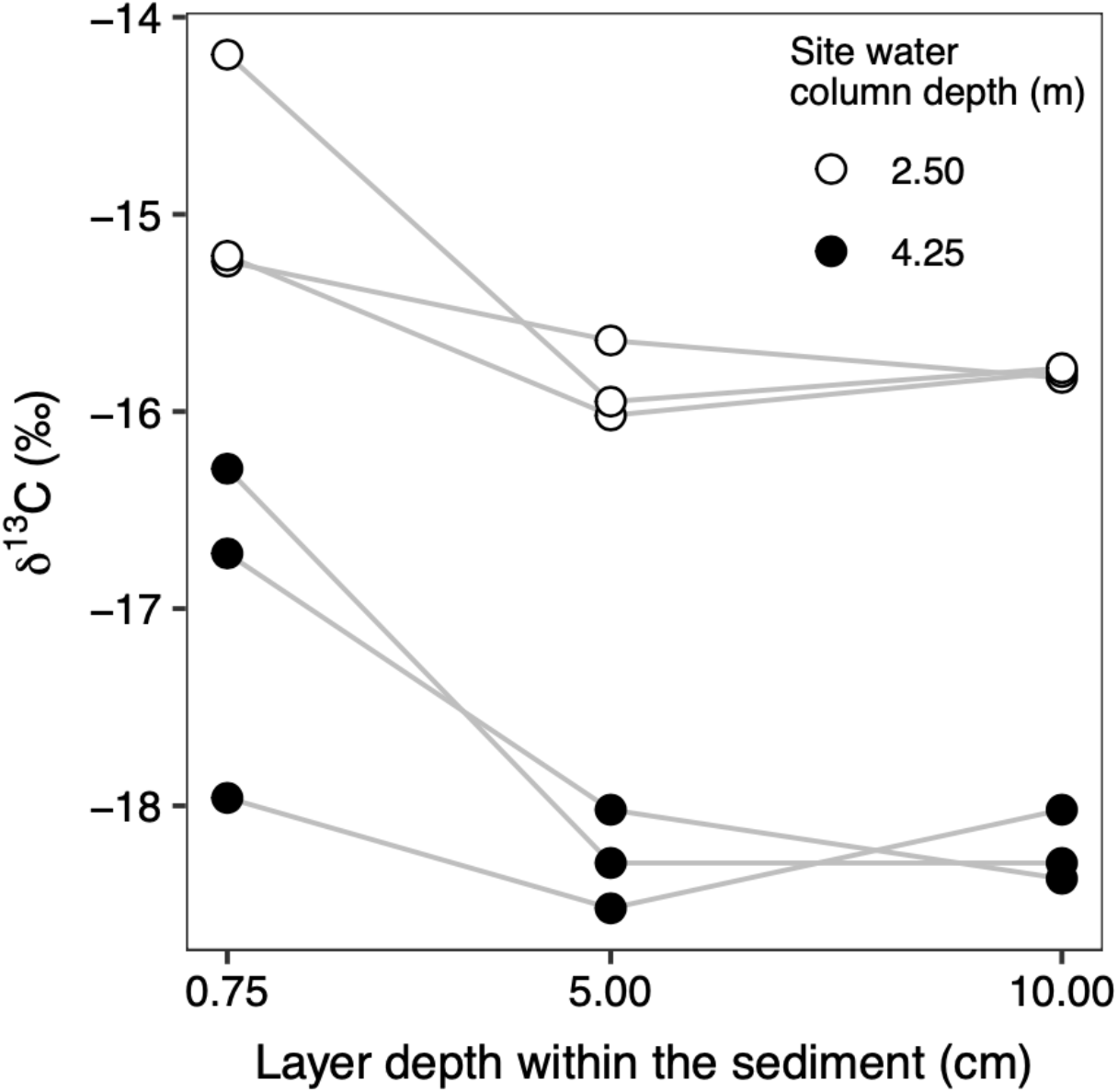
Sediment δ^13^C signatures from multiple layer depths within cores collected from two sites with water column depths of 2.50 and 4.25 m. Within each core, sediment layers were sampled at 0.75 cm (i.e., the surface layer), 5 cm, and 10 cm below the sediment’s surface. Each point shows a sample, with lines connecting layers from an individual core.

## Discussion

In this study, we examined patterns of larval abundance and resource use for two midge species that dominate the benthos of Lake Mývatn. Consumer-resource interactions likely drive the high-amplitude fluctuations of the population of *T. gracilentus* in Mývatn (Einarsson et al. 2002), and population crashes of five orders-of-magnitude suggest that benthic resources can be limiting, at least intermittently. Moreover, the synchrony in *T. gracilentus* and *C. islandicus* population fluctuations (Gardarsson et al., 2004) suggests that both species may be limited by the same resources. Together, the ecological similarity of *T. gracilentus* and *C. islandicus*, their high peak population abundances, and their synchronized fluctuations at the whole-lake level warrant investigating possible factors contributing to their coexistence.

Habitat partitioning was considered as a possible explanation for the coexistence of *T. gracilentus* and *C. islandicus*. Based on the positive correlation in their abundances across sediment cores (20 cm^2^), this form of spatial partitioning does not appear to be strong. In supplemental analyses, positive correlations in abundance were also found at the same site (several m^2^) sampled in the same day and at sites spanning the lake’s main basin within a given year, although these were fairly moderate associations (Supporting Information; Fig. S2). Altogether, the lack of negative correlations in larval abundances implies that *T. gracilentus* and *C. islandicus* generally occur within the same areas rather than occupying distinct areas across the lake, at least over the duration of this study that occurred during time periods of relatively high population abundances of both species when competition is likely to be important. Their positive correlation in abundance may be due to shared responses to resource abundance or local abiotic conditions, but it does not necessarily exclude the possibility that *T. gracilentus* and *C. islandicus* compete with one another. Negative effects from interspecific competition could still occur, and in the absence of interspecific competition, associations between *T. gracilentus* and *C. islandicus* could have been even more positive. Nonetheless, these patterns suggest that any potential negative effects of interspecific interactions at these spatial scales are overridden by factors leading to positive correlations in larval abundances.

We assessed partitioning of food resources between *T. gracilentus* and *C. islandicus* with stable carbon isotope values. The interspecific differences in δ^13^C signatures suggest differences in resource use. We cannot definitively infer resource partitioning between Mývatn’s *T. gracilentus* and *C. islandicus* populations from these results, because we cannot demonstrate that differences in resource use decrease interspecific competition (Colwell and Futuyma 1971). Nonetheless, our results illustrate the *potential* for resource partitioning between these two ecologically and dynamically similar populations. These resource use differences may have occurred due to one or more mechanisms: (i) finer horizontal spatial partitioning (< 20 cm^2^) than captured by our sediment core sampling, (ii) fine-scale differences in resource selectivity, and (iii) tube-building behaviors associated with contrasting feeding styles and vertical distributions within the sediment. These mechanisms are not mutually exclusive and are discussed together below.

It is reasonable to consider surface sediment as the putative resource for *T. gracilentus* and *C. islandicus*, and both species likely acquire some portion of their diet from feeding within this habitat. However, such feeding could occur with contrasting resource selectivity at fine spatial scales, as indicated by our stable isotope data. Specifically, *T. gracilentus* δ^13^C values were higher compared to surface sediment, while *C. islandicus* values were lower. Furthermore, the degrees of differentiation between larvae and surface sediment—mean difference in δ^13^C is 1.84‰ for *T. gracilentus* and –2.87‰ for *C. islandicus*—are greater than expected due to trophic fractionation alone and suggest discriminate feeding behavior. For example, previous estimates of ^13^C-trophic fractionation include 0.3‰ from a chironomid feeding study (Doi et al. 2006) and −0.4‰ for aquatic herbivores from a meta-analysis (Vander Zanden and Rasmussen 2001).

The degree of enrichment in *T. gracilentus* δ^13^C values compared to surface sediments provides insight about their feeding and indicates that they selectively feed on ^13^C-enriched resources. Elevated algal δ^13^C signatures are associated with high rates of primary production, because increased photosynthetic rates can induce inorganic carbon limitation and lead to lower discrimination against the heavy ^13^C isotope during carbon fixation (Hecky and Hesslein 1995, Hill and Middleton 2006, Devlin et al. 2013). Our surface sediment samples contained the top 0.75-cm layer of sediment; however, the most productive algal biofilm is likely restricted within the uppermost part of this layer and likely has comparatively enriched δ^13^C values. *T. gracilentus* larvae may selectively consume the most productive algal material within the photosynthetically active layer (e.g., rapidly growing diatom cells). This explanation was proposed by Devlin et al. (2013) to explain the high δ^13^C values of benthic grazers relative to surface sediment. Given the smaller body size of *T. gracilentus* larvae compared to *C. islandicus*, it is also possible that *T. gracilentus* feed on smaller, more rapidly growing diatom species, which may be associated with lower ^13^C-discrimination (Korb et al. 1996).

The low *C. islandicus* isotope signatures compared to surface sediment suggests selectivity for certain ^13^C-depleted resources (Solomon et al., 2008), including detritus and/or microbes that assimilate detrital carbon (Fiskal et al. 2021). One possibility is that *C. islandicus* may more heavily rely on microbes that process organic matter at the sediment surface. Aerobic heterotrophic bacteria discriminate against ^13^C, especially when carbon is non-limiting (McGoldrick et al. 2008). Surface sediments at Mývatn have relatively high organic matter content (23 ± 3%) (Ives et al. 2021), such that detritus-associated bacteria are unlikely to be carbon-limited and are potentially ^13^C-depleted (McGoldrick et al. 2008). Selectivity for such microbes can affect consumer δ^13^C values; for example, low δ^13^C values of *Chironomus tentans* relative their detrital food source were attributed to the assimilation of microbial biomass growing on the detritus (McGoldrick et al. 2008). Other work has found that microbial communities of chironomids are dominated by carbohydrate-degrading bacteria, similarly suggesting the potential for larvae to preferentially rely on resources linked to algal detritus in surface sediments (Fiskal et al. 2021).

An additional explanation for the low *C. islandicus* δ^13^C values relative to *T. gracilentus* is that *C. islandicus* selectively feeds on pelagic-sourced carbon. Previous studies have reported that other *Chironomus* species consume phytoplankton or recently deposited phytodetritus from the sediment’s surface (i.e., deposit-feeding) (Johnson 1985, 1987, Gullberg et al. 1997, Doi et al. 2006). Such feeding habits can affect consumer isotope signatures because pelagic-derived carbon resources are typically ^13^C-depleted relative to those from benthic habitats (Hecky and Hesslein 1995). However, our ability to assess the reliance of *C. islandicus* on pelagic-derived carbon is limited because differences in δ^13^C values between paired surface sediment and pelagic samples from Mývatn are low (mean difference ± SD = 2.65 ± 1.71‰; Supporting Information; Fig. S3a) compared to other studies (7.6‰ in Doi et al. 2010; 11.7-15.3‰ in Chételat et al. 2010). Moreover, *C. islandicus* δ^13^C signatures relative to those of pelagic samples collected from the same site at around the same time are variable, including the direction of relative enrichment (difference = –0.69 ± 2.09‰; Supporting Information; Fig. S3b). Thus, we cannot provide conclusive evidence for or against the use of pelagic-derived carbon by *C. islandicus*.

Beyond the potential for contrasting resource selectivity at the sediment surface, an additional, nonexclusive explanation for differences in resource use between *T. gracilentus* and *C. islandicus* involves their respective tube-building behaviors that are associated with contrasting feeding styles and vertical distributions within the sediment. Several species in the genus *Tanytarsus*, including *T. gracilentus*, build vertical tubes that at high density are arranged against one another and extend above the sediment surface, such that larvae generally remain in the top 2 cm of sediment (Heinis et al. 1994, Chaloner and Wotton 1996, Ólafsson and Paterson 2004). Furthermore, *T. gracilentus* tubes provide high-quality substrate for benthic algae at the sediment surface, thereby increasing benthic chl-a concentrations and primary production (Phillips et al. 2019). If *T. gracilentus* rely heavily on productive diatoms that they ‘garden’ from their tubes (sensu Ings et al. 2012), it could facilitate their selection of ^13^C-enriched benthic resources, thereby elevating their δ^13^C values above the bulk surface sediment.

In contrast, *C. islandicus* create “blind-ended” or “I-shaped” (Kristensen et al. 2012) burrows that extend 10-15 cm below the sediment surface in Mývatn (Herren et al. 2017), and other studies have similarly documented *Chironomus* spp. at sediment depths of 7-12 cm (Heinis et al. 1994, Charboneau and Hare 1998). Ventilation activity pumps water and particulate organic matter into *Chironomus* spp. burrows, such that larvae effectively filter particles that become entrapped within their tubes (Walshe 1947, Kristensen et al. 2012, Holker et al. 2015). Furthermore, observations of *C. islandicus* in laboratory microcosms reveal that they actively feed upon the inner lining of their tubes below the sediment surface (Ólafsson, unpublished data). Distinctions between *C. islandicus* and surface sediment δ^13^C values could be explained if the particles that enter their burrows are relatively ^13^C-depleted. For example, if productive diatom biofilms, with corresponding enriched δ^13^C values, are largely attached to *T. gracilentus* tubes, they may be less likely to be transported into *C. islandicus* burrows. Rather, ventilation currents may more likely transport unattached detrital particles at the sediment-water interface or suspended phytoplankton. Altogether, correspondence between *T. gracilentus* and *C. islandicus* feeding activities and the distributions of their tubes may facilitate resource partitioning vertically within the sediment habitat.

*C. islandicus* δ^13^C values could also be affected by the isotope signatures of organic matter below the sediment surface, as sediment layers at 5 cm and 10 cm depth were ^13^C-depleted compared to the surface layer. The isotope signatures of deeper sediment layers are not necessarily indicative of resources attached to the inner lining of *C. islandicus* burrows because many of these particles likely come from the sediment-water interface. However, *C. islandicus* may include material from deeper within the sediment profile when constructing their tubes, thereby potentially incorporating relatively ^13^C-depleted particles within the inner burrow wall upon which they graze. Additionally, previous research has demonstrated that multiple species of *Chironomus* feed upon older particles from below the oxic-anoxic sediment interface (e.g., underlying the oxygenated environment of their burrows) (Martin et al. 2008, Proulx and Hare 2014). While there have been no direct observations of *C. islandicus* feeding beyond their tube lining, such behavior would likely result in the consumption of ^13^C-depleted organic matter. The depleted δ^13^C signatures of organic matter 5-10 cm below the surface may be due to detrital processing within the sediment, whereby selective degradation of particular organic matter compounds can alter the isotopic signature of the remaining organic matter pool (Meyers and Ishiwatari 1993, Lehmann et al. 2002, Canuel and Hardison 2016). For example, δ^13^C depletion in deeper sediment layers may be explained by the more-rapid degradation of labile, ^13^C-enriched compounds (e.g., carbohydrates, proteins), which leads to the retention of specific recalcitrant organic matter fractions that are comparatively ^13^C-depleted (Meyers and Ishiwatari 1993, Lehmann et al. 2002). Overall, if *C. islandicus* spend ample time 5-10 cm below the sediment surface, they likely encounter organic matter pools with relatively low isotopic signatures.

Finally, consumption of microbes involved in methane cycling, such as methane oxidizing bacteria, is a well-documented contributor to low δ^13^C signatures of chironomids living within anoxic sediment environments (Jones et al. 2008). Methane-oxidizing bacteria can have very depleted δ^13^C values due to fractionation during biogenic methane production and additional methanotrophic discrimination against ^13^C (Kiyashko et al. 2001, Grey et al. 2004, Hershey et al. 2006, Jones et al. 2008). However, our *C. islandicus* isotope signatures are not near the levels of ^13^C-depletion as observed in consumers with substantial dietary contributions from methane-derived carbon (Kiyashko et al. 2001, Grey et al. 2004, Jones et al. 2008, Proulx and Hare 2014, Fiskal et al. 2021). Moreover, dissolved oxygen concentrations near the sediment surface generally remain high throughout the summer in Mývatn, while substantial reliance of chironomid larvae on methane-derived carbon seems restricted to lakes with late-summer hypoxia (< 2mg O_2_/L) at the sediment-water interface (Jones et al 2008). Thus, we suggest that methane-derived carbon is unlikely to be an important contributor to the diet of *C. islandicus* in Mývatn, although even minor assimilation of such an extremely ^13^C-depleted resource could decrease consumer signatures (Solomon et al. 2008).

Despite the interspecific differences in δ^13^C values, both *T. gracilentus* and *C. islandicus* larval δ^13^C values decreased across a water-column depth gradient. This suggests that δ^13^C of both consumers in part reflects the isotopic characteristics of the sediment, as surface sediment δ^13^C also decreased with depth. The declines in δ^13^C values of *T. gracilentus*, *C. islandicus* and surface sediment may reflect decreasing primary productivity at deeper depths of the lake (Devlin et al. 2013). This pattern would be consistent with our interpretation that high δ^13^C values of *T. gracilentus* reflect their use of productive diatoms on their tubes, since deeper sites generally have lower light levels and hence lower algal growth (Phillips et al. 2019). However, other studies have attributed the negative relationship between depth and δ^13^C values to increased phytoplankton deposition (Karlsson and Byström 2005, Doi et al. 2006) or increased prevalence of organic matter emanated from respired carbon and methanogenesis (Solomon et al. 2008).

In addition to differences in resource use, it is worth considering whether differences in larval development time between *C. islandicus* and *T. gracilentus* affected comparisons of their δ^13^C values. Isotope signatures represent a time-integrated account of consumer assimilation, with a time lag before a consumer’s isotopic signature reaches equilibrium with its diet (Vander Zanden et al. 2015). Therefore, a consumer’s δ^13^C value may not match its current diet if it switches resources or if its resource has temporally variable δ^13^C signatures (Forbes and Gratton 2011). This raises the possibility that the longer larval life stage of *C. islandicus* may mean that their isotopic signatures integrate resource δ^13^C values over periods that *T. gracilentus* signatures do not encompass. For example, this would be the case if *C. islandicus* larval isotope signatures in summer retained resource signatures from the preceding winter, whereas *T. gracilentus* signatures of summer generation larvae would only reflect resource signatures since the emergence in June. However, this possibility is unlikely given the short expected δ^13^C half-life for both species; based on quantitative relationships reported within Thomas and Crowther (2015) and Vander Zanden et al. (2015), and individual body masses reported from Herren et al. (2017), the δ^13^C half-life of both taxa will be on the order of days, rather than weeks or months. Along these lines, in a reciprocal transplant experiment, δ^13^C values of *Chironomus* and *Stictochironomus* larvae rapidly shifted in response to new sediment conditions (Hershey et al. 2006), suggesting that chironomid larvae equilibrate to isotopic shifts of available resources relatively quickly.

## Conclusions

While competitive interactions between species can cause negative correlations in their abundances, competitors can show positive correlations in time and space if they rely on resources that themselves fluctuate. Thus, the positive correlations in *T. gracilentus* and *C. islandicus* abundance across sites in Mývatn and previously documented synchronous population dynamics at the whole-lake level (Gardarsson et al., 2004) imply that *T. gracilentus* and *C. islandicus* use at least some of the same resources, or different resources that themselves show positive correlations. Given their similar proximity to sediment resources and previous diet studies (Ólafsson 1987 as reported within Einarsson et al. 2004; Ingvason 2002, Ingvason et al. 2002, 2004), *T. gracilentus* and *C. islandicus* almost certainly consume similar resources (e.g., diatoms, detritus, microbes) and could possibly compete for at least some of them. Even though differences in carbon isotopic signatures do not imply complete non-overlap in diets, we found that *T. gracilentus* and *C. islandicus* feed on sufficiently different resources to give distinct δ^13^C signatures. These consistent isotopic differences suggest that a degree of resource partitioning is likely. Specifically, we propose that *T. gracilentus* predominantly acts as a grazer, feeding selectively on productive algae from their tubes, while *C. islandicus* relies more heavily on detrital-associated resources that they collect from the sediment surface and their sub-surface burrows. These resources may themselves be dynamically coupled and related to the synchronous dynamics of the consumer species. For example, algal dynamics at the benthic surface might be coupled to fluctuations in organic matter availability or heterotrophic activity further down in the sediment. While the factors explaining the synchrony of population fluctuations of *T. gracilentus* and *C. islandicus* remain uncertain, resource partitioning might explain both how they coexist and how they can each simultaneously achieve remarkably high densities. Resource use differences at fine spatial scales may be similarly important for understanding patterns of abundance for other ecologically similar species.

## Acknowledgements

Árni Einarsson provided invaluable assistance in setting up the long-term monitoring, ongoing logistical support at the Mývatn Research Station, and helpful comments on this manuscript. We also thank the many interns who assisted with sample collection in the field and sorting of larvae and C. Gratton for guidance and logistical support in preparing samples for isotopic analysis.

## Supporting Information

### Supplemental analysis

#### Correlation in larval abundance

Our correlation analysis presented in the main text uses species-specific larval abundances from individual sediment cores. To supplement this analysis, we examined correlations at two higher scales: (i) aggregated across site-date combinations (i.e., using mean species-specific abundances for each site-date combination), and (ii) aggregated across site-year combinations (i.e., using mean species-specific abundances for site-year combinations). When calculating the correlation between abundances aggregated at the site-date level, we standardized abundances for each species by subtracting the species-specific mean abundance for each site-year combination. We report Pearson’s correlation coefficient for these analyses.

Abundances of *T. gracilentus* and *C. islandicus* larvae were positively correlated at the levels of aggregated site-date combinations (*r* = 0.43; Fig. S2a) and aggregated site-year combinations (*r* = 0.51; Fig. S2b).

#### Pelagic δ^13^C signatures

During the study period, we collected a limited number of pelagic samples and submitted them for isotopic analysis. For these samples, we collected field water with a modified Schindler trap at approximately 1-m intervals. We avoided sampling within 1 m of the sediment surface to prevent disrupting the sediment. A total of 10-30 L of water per sample was homogenized and filtered through a 63-µm sieve in the field. Contents suspended in the water column that were retained on the 63-µm sieve included phytoplankton, zooplankton, and other particulate organic matter. Pelagic isotope samples were then transferred to glass vials and dried at 60 °C. Once dry, the samples were processed following the same procedures as described in the main text for larvae and surface sediment (**Methods:** *Stable isotope samples*).

In the main text, we discuss how reliance of pelagic-derived carbon may influence *C. islandicus* larvae δ^13^C signatures (**Discussion**). To supplement this discussion, we present a visualization of δ^13^C values from pelagic samples compared to those of (i) surface sediment and (ii) *C. islandicus* larvae (Fig. S3). For this visualization, we included a subset of our surface sediment and *C. islandicus* samples that had a corresponding pelagic δ^13^C value. Like the main text (**Methods:** *Statistical analysis*), samples that were paired from a given site were collected on the same date in most cases, but for some pairs, we compared samples that were collected 1-12 days apart.

**Figure S1.**
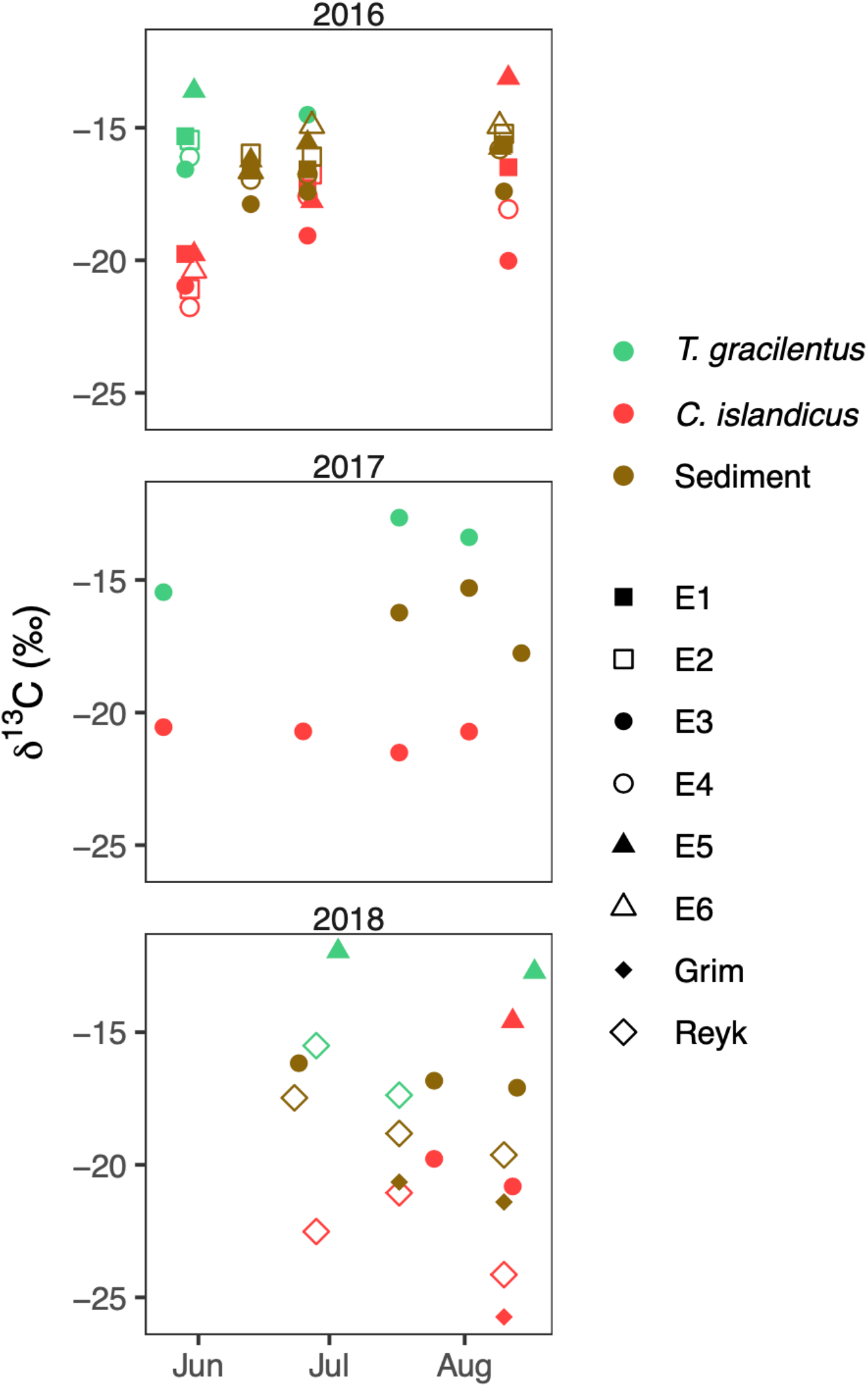
δ^13^C signatures of *T. gracilentus* larvae, *C. islandicus* larvae, and surface sediments collected from sites in Lake Mývatn during 2016-2018.

**Figure S2.**
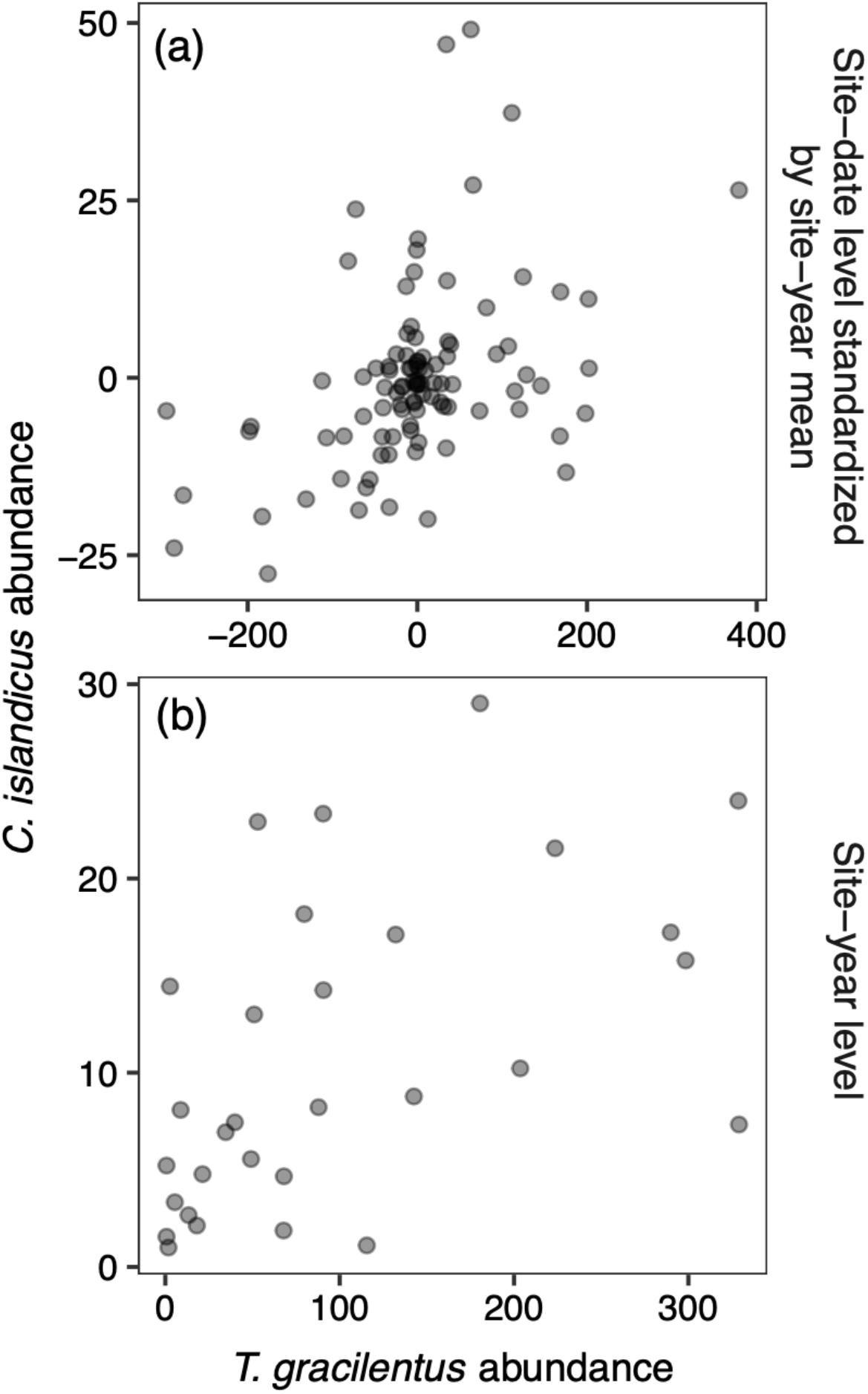
Comparisons of *T. gracilentus* and *C. islandicus* abundances at the levels of (a) larvae aggregated for site-date combinations, and (b) larvae aggregated for site-year combinations. In panel (a), abundances were standardized by subtracting the mean species-specific abundance for each site-year combination.

**Figure S3.**
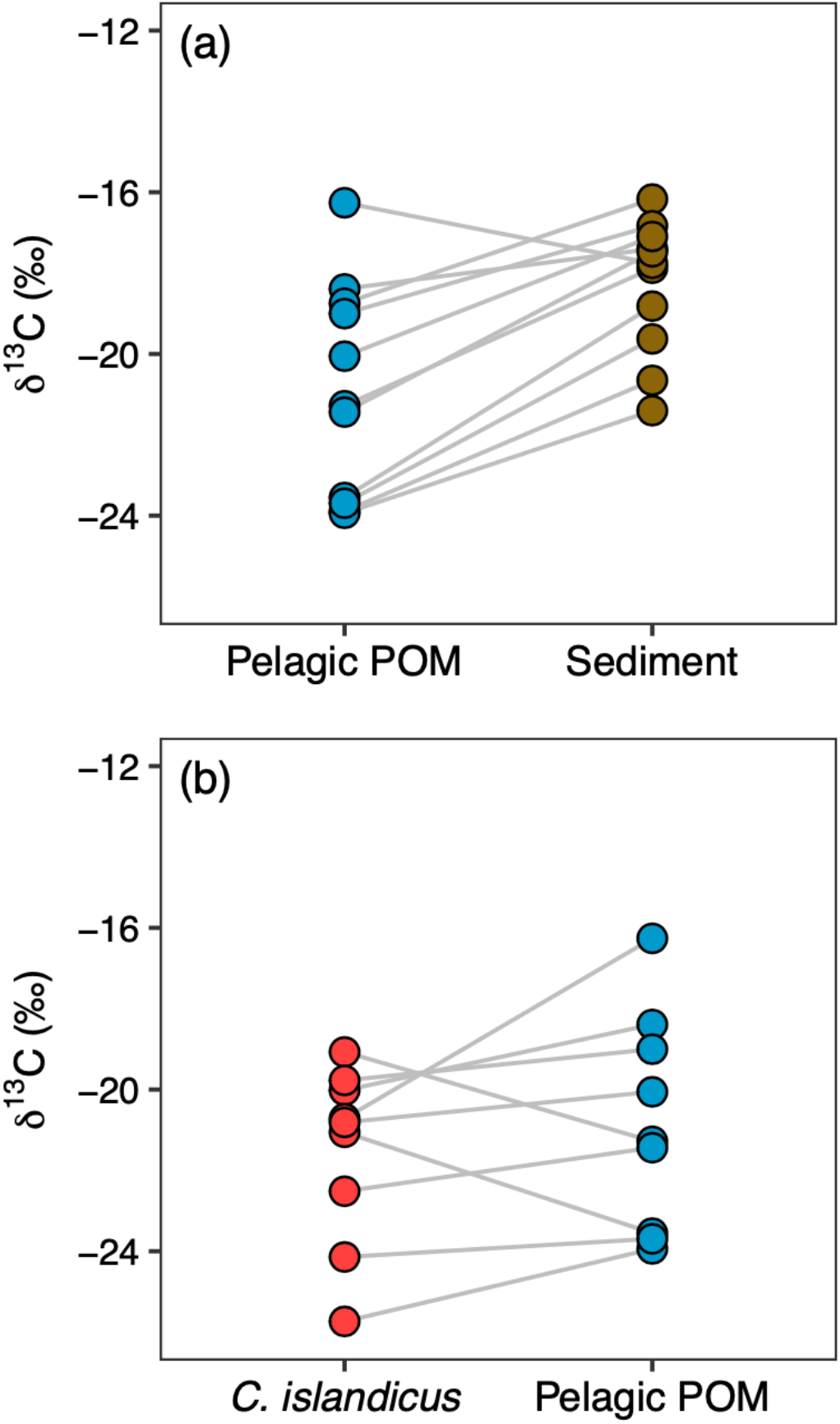
Comparisons of δ^13^C signatures among (a) pelagic samples and surface sediment and (b) *C. islandicus* larvae and pelagic samples. Lines connect a pair of samples collected from the same site and near the same date.

**Table S1.** List of larvae and sediment samples collected for stable isotope analysis from 2016-2018.

| Sample type | Sample date | Site depth (m) | Site name | n | $\delta^{13}\text{C}$ (SD) |
| --- | --- | --- | --- | --- | --- |
| *C. islandicus* | 5/29/2016 | 2.7 | E1 | 1 | -19.76 |
| *C. islandicus* | 5/29/2016 | 3.3 | E3 | 1 | -20.97 |
| *C. islandicus* | 5/30/2016 | 2.7 | E2 | 1 | -21.08 |
| *C. islandicus* | 5/30/2016 | 2.9 | E4 | 1 | -21.77 |
| *C. islandicus* | 5/31/2016 | 2.5 | E5 | 1 | -19.77 |
| *C. islandicus* | 5/31/2016 | 2.8 | E6 | 1 | -20.39 |
| *C. islandicus* | 6/26/2016 | 2.7 | E1 | 1 | -17.27 |
| *C. islandicus* | 6/26/2016 | 3.3 | E3 | 1 | -19.07 |
| *C. islandicus* | 6/26/2016 | 2.9 | E4 | 1 | -17.58 |
| *C. islandicus* | 6/27/2016 | 2.7 | E2 | 1 | -16.76 |
| *C. islandicus* | 6/27/2016 | 2.5 | E5 | 1 | -17.77 |
| *C. islandicus* | 8/11/2016 | 2.7 | E1 | 1 | -16.49 |
| *C. islandicus* | 8/11/2016 | 3.3 | E3 | 1 | -20.02 |
| *C. islandicus* | 8/11/2016 | 2.9 | E4 | 1 | -18.07 |
| *C. islandicus* | 8/11/2016 | 2.5 | E5 | 1 | -13.12 |
| *C. islandicus* | 5/24/2017 | 3.3 | E3 | 1 | -20.55 |
| *C. islandicus* | 6/25/2017 | 3.3 | E3 | 1 | -20.71 |
| *C. islandicus* | 7/17/2017 | 3.3 | E3 | 1 | -21.51 |
| *C. islandicus* | 8/2/2017 | 3.3 | E3 | 1 | -20.72 |
| *C. islandicus* | 6/28/2018 | 4.25 | Reyk | 4 | -22.52 (0.26) |
| *C. islandicus* | 7/17/2018 | 4.25 | Reyk | 4 | -21.06 (1.35) |
| *C. islandicus* | 7/25/2018 | 3.3 | E3 | 3 | -19.77 (1.68) |
| *C. islandicus* | 8/10/2018 | 5.1 | Grim | 1 | -25.74 |
| *C. islandicus* | 8/10/2018 | 4.25 | Reyk | 4 | -24.14 (1.09) |
| *C. islandicus* | 8/12/2018 | 3.3 | E3 | 2 | -20.81 (0.82) |
| *C. islandicus* | 8/12/2018 | 2.5 | E5 | 3 | -14.58 (1.06) |
| *T. gracilentus* | 5/29/2016 | 2.7 | E1 | 1 | -15.32 |
| *T. gracilentus* | 5/29/2016 | 3.3 | E3 | 1 | -16.57 |
| *T. gracilentus* | 5/30/2016 | 2.7 | E2 | 1 | -15.46 |
| *T. gracilentus* | 5/30/2016 | 2.9 | E4 | 1 | -16.10 |
| *T. gracilentus* | 5/31/2016 | 2.5 | E5 | 1 | -13.61 |
| *T. gracilentus* | 6/26/2016 | 3.3 | E3 | 1 | -14.51 |
| *T. gracilentus* | 5/24/2017 | 3.3 | E3 | 1 | -15.46 |
| *T. gracilentus* | 7/17/2017 | 3.3 | E3 | 1 | -12.65 |
| *T. gracilentus* | 8/2/2017 | 3.3 | E3 | 1 | -13.39 |
| *T. gracilentus* | 6/28/2018 | 4.25 | Reyk | 3 | -15.51 (0.16) |
| *T. gracilentus* | 7/3/2018 | 2.5 | E5 | 3 | -11.95 (0.15) |
| *T. gracilentus* | 7/17/2018 | 4.25 | Reyk | 4 | -17.37 (0.09) |
| *T. gracilentus* | 8/17/2018 | 2.5 | E5 | 3 | -12.72 (0.10) |
| Surface sediment | 6/13/2016 | 2.7 | E1 | 1 | -16.59 |
| Surface sediment | 6/13/2016 | 2.7 | E2 | 1 | -15.98 |
| Surface sediment | 6/13/2016 | 3.3 | E3 | 1 | -17.88 |
| Surface sediment | 6/13/2016 | 2.9 | E4 | 1 | -16.96 |
| Surface sediment | 6/13/2016 | 2.5 | E5 | 1 | -16.23 |
| Surface sediment | 6/13/2016 | 2.8 | E6 | 1 | -16.64 |
| Surface sediment | 6/26/2016 | 2.7 | E1 | 1 | -16.57 |
| Surface sediment | 6/26/2016 | 3.3 | E3 | 1 | -17.42 |
| Surface sediment | 6/26/2016 | 2.9 | E4 | 1 | -16.76 |
| Surface sediment | 6/26/2016 | 2.5 | E5 | 1 | -15.56 |
| Surface sediment | 6/27/2016 | 2.7 | E2 | 1 | -16.09 |
| Surface sediment | 6/27/2016 | 2.8 | E6 | 1 | -14.93 |
| Surface sediment | 8/9/2016 | 2.9 | E4 | 1 | -15.81 |
| Surface sediment | 8/9/2016 | 2.5 | E5 | 1 | -15.76 |
| Surface sediment | 8/9/2016 | 2.8 | E6 | 1 | -14.94 |
| Surface sediment | 8/10/2016 | 2.7 | E1 | 1 | -15.63 |
| Surface sediment | 8/10/2016 | 2.7 | E2 | 1 | -15.23 |
| Surface sediment | 8/10/2016 | 3.3 | E3 | 1 | -17.40 |
| Surface sediment | 7/17/2017 | 3.3 | E3 | 1 | -16.23 |
| Surface sediment | 8/2/2017 | 3.3 | E3 | 1 | -15.30 |
| Surface sediment | 8/14/2017 | 3.3 | E3 | 1 | -17.76 |
| Surface sediment | 6/23/2018 | 4.25 | Reyk | 1 | -17.47 |
| Surface sediment | 6/24/2018 | 3.3 | E3 | 1 | -16.17 |
| Surface sediment | 7/17/2018 | 5.1 | Grim | 1 | -20.65 |
| Surface sediment | 7/17/2018 | 4.25 | Reyk | 1 | -18.82 |
| Surface sediment | 7/25/2018 | 3.3 | E3 | 1 | -16.83 |
| Surface sediment | 8/10/2018 | 5.1 | Grim | 1 | -21.40 |
| Surface sediment | 8/10/2018 | 4.25 | Reyk | 1 | -19.63 |
| Surface sediment | 8/13/2018 | 3.3 | E3 | 1 | -17.09 |

